# Expression of nitric oxide synthase and nitric oxide levels in peripheral blood cells and oxidized low-density lipoprotein levels in saliva as early markers of severe dengue

**DOI:** 10.1101/664151

**Authors:** Harsha Hapugaswatta, Ruwani L. Wimalasekara, Suharshi S. Perera, Ranjan Premaratna, Kapila N. Seneviratne, Nimanthi Jayathilaka

## Abstract

**Background:** Severe dengue (SD), experienced by only a fraction of dengue patients can be lethal. Due to lack of early markers that can predict the evolution of SD, all dengue patients have to be monitored under hospital care. We discovered early oxidative stress markers of SD to identify patients who can benefit from early intervention before the symptoms appear.

**Methods:** Expression of inducible nitric oxide synthase (iNOS) in peripheral blood cells (PBC), nitric oxide (NO) and oxidized low-density lipoprotein (oxLDL) levels in plasma and saliva collected at early stages of dengue infection from 20 non-severe dengue fever (DF) patients and 20 patients who later developed SD were analyzed in a retrospective nested case-control study.

**Results:** Expression of iNOS is significantly (P<0.05) lower in patients who developed SD than in DF patients at admission within 4 days from fever onset. Median plasma NO concentration within 4 days from fever onset is also significantly (P<0.05) lower in patients who developed SD (17.9±1.6 μM) than DF (23.0±2.1 μM). Median oxLDL levels in plasma within 3 days from fever onset is significantly (P<0.05) lower in patients who developed SD (509.4±224.1 ng/mL) than DF (740.0±300.0 ng/mL). Median salivary oxLDL levels are also significantly (P<0.05) lower in patients who developed SD (0.8±0.5 ng/mL) than DF (3.6±2.6 ng/mL) within 4 days from fever onset.

**Conclusions:** These findings suggest that the expression of iNOS (73% sensitivity, 86% specificity) and plasma NO (96% sensitivity, 61% specificity at 22.3 μM; P<0.05) may serve as early markers of SD within 3 days from fever onset. Salivary oxLDL levels may serve as early non-invasive markers of SD with a sensitivity and specificity respectively of 57% and 91% at 0.9 ng/mL, 76% and 55% at 2.3 ng/mL and 100% and 50% at 4.6 ng/mL; P<0.05) within 4 days from fever onset.

## 1. Introduction

Dengue is an extremely prevalent mosquito-borne viral disease in many tropical countries including Sri Lanka. It is the second most important tropical disease (after malaria) with 284 – 528 million dengue infections resulting in 67 – 136 million clinically manifested dengue cases with half the global population at-risk posing a significant public health threat worldwide [1, 2] Currently, the number of SD cases in Sri Lanka has dramatically increased. In 2017, 1,86,101 suspected dengue cases were reported to the Epidemiology Unit of Sri Lanka from all over the island while 2018 only saw 51659 reported cases of dengue. However, 104667 cases were reported in 2019 [3]. Most people infected with dengue viruses are asymptomatic while others may suffer a wide range of clinical manifestations from mild fever to SD. DF is a serious, debilitating condition and severe manifestations of the disease are major causes of hospitalization and death, globally [1, 4]. Unlike DF, SD is characterized by severe possibly lethal vasculopathy marked with plasma leakage, intrinsic coagulopathy and massive internal bleeding [5, 6].

Despite the social and clinical impact, there are no antiviral therapies available for treatment of dengue [7]. The vaccine that is licensed in 18 countries has several limitations because it is only recommended for individuals aged 9-45 years who have had previous exposure to dengue [8]. As such, prevention is mostly limited to vector control measures. Several efficient and relatively reliable diagnostic tests based on PCR or serological testing are available for detection of dengue virus infections. Rapid lateral flow tests for the presence of dengue NS1 antigen are most commonly administered for dengue diagnosis in Sri Lanka. These diagnostic tests, however, do not distinguish between DF and SD [9]. Limited progress has been made in finding markers that can indicate the evolution of dengue infection to the severe form of the illness at an early stage of infection. Infact, a diagnosis of disease severity is usually made after the patient is presented with SD symptoms. Several studies have compared the transcriptomes of patients that developed DF with those who developed SD to identify molecular markers such as cytokines associated with disease severity [6, 10, 11–16]. In a genome-wide association study, genetic varients in major histocompatibility complex class I polypeptide-related sequence B gene (MICB) and phospholipase C epsilon 1 (PLCE1) has been found to be associated with SD [17]. A later study revealed that these genetic varients are not only associated with SD but also with less severe clinical phenotypes of DF [18]. Allelic forms of MICA and MICB, on the other hand, has been found to strongly associate with susceptibility to illness but not with the severity of infection [19]. Recent studies have also reported differential expression of microRNA in dengue patients and in infected cultured cells [20–24]. However, most of the patient studies do not limit the sample pool to acute phase of infection at which differentail diagnosis is not possible. Thus, despite these efforts, endeavors to discover a prognostic test for SD is yet to become part of the dengue clinical tests, leaving much to be done in finding a solution to this public health crisis.

Inducible nitric oxide synthase (iNOS) has been implicated in host response to dengue virus infection [25]. Expression of iNOS leads to nitric oxide (NO) biosynthesis resulting in generation of a highly reactive nitrogen oxide species, peroxynitrite, via a radical coupling reaction of NO with superoxide which in turn causes potent oxidation and nitration reactions of various biomolecules including lipids. NO and oxygen radicals such as superoxide are key molecules in the pathogenesis of various infectious diseases. [26]. NO biosynthesis through expression of iNOS occurs in a variety of microbial infections. iNOS mediated production of nitric oxide, accumulation of reactive oxygen species (ROS) and reactive nitrogen species (RNS), as well as perturbation of the levels of the intracellular antioxidant, glutathione (GSH) have been reported in DENV infected human cells and animal models. Elevated lipid and protein oxidation markers in serum and plasma and alteration in the redox status of DENV-infected patients have been associated with increased inflammatory responses, cell death, increased susceptibility to DENV infection and increased disease severity [27]. iNOS activity and plasma NO has been implicated in inflammatory responses and plasma leakage [25]. SD is characterized by thrombocytopenia (low platelet count) and plasma leakage. Therefore, NO which plays a complex and diverse physiological and pathophysiological role may serve as an early prognostic marker in dengue [28]. A recent study indicates the potential of the levels of serum NO in DF and SD patients as an early marker of disease diagnosis [29]. However, the levels of NO were not evaluated for their potential as early markers of the disease in other biological fluids. Further, differential expression of iNOS during the early stages of DENV infection among patients who do not present severe symptoms and those who develop severe symptoms has not been evaluated to assess the potential to serve as an early marker of disease severity. *In vitro* studies of DENV and other flavivirus infections such as Japanese encephalitis virus and West Nile virus suggest that lipids and lipoproteins may play a role in modifying virus infectivity of target cells due to the role of cholesterol-rich lipid rafts in flavivirus entry. In fact, lower total serum cholesterol and low-density lipoprotein cholesterol levels have been shown to associate with subsequent risk of developing severe symptoms of dengue such as dengue hemorrhagic fever/dengue shock syndrome using the WHO 1997 dengue severity classification [30]. Therefore, ROS mediated oxidation of low-density lipoprotein cholesterol levels may also serve as an early marker of disease severity in dengue. Studies have not been conducted to evaluate the effect of severity of DENV infection on the levels of oxidized low-density lipoprotein during the early stages of infection when differential prognosis is not possible. Therefore, in this study, we evaluated whether the severity of dengue infection is correlated with early differential expression of iNOS and resultant changes in NO levels and oxLDL levels in plasma from patients who tested positive for dengue within 4 days from fever onset before severe symptoms are presented. Levels of biomarkers such as LDL in saliva has been reported to correlate with serum and plasma LDL levels [31]. Since saliva is a safe and easy to handle biological fluid that can be collected using non-invasive measures, we also evaluated the salivary oxLDL levels in samples from patients presented with symptoms of dengue fever during the early stages of infection compared to those who later developed SD.

The development and the severity of SD can be mitigated with proper disease management. If diagnosed early, before severe symptoms are presented, effective disease management of SD only involves hospital care and hydration. Identifying early molecular markers of SD may help distinguish dengue patients who would benefit from early intensive therapy and hospitalization before severe symptoms appear and increase the availability of public health resources and also mitigate the cost of public health and the impact on the national economy.

## 2. Materials and Methods

### 2.1. Sample collection and processing

127 adult male and female patients (above age 18) presented with clinical symptoms of dengue viral infection according to WHO Dengue case classification (2009) (fever, with two of the following criteria: nausea, vomiting, rash, aches and pains, positive tourniquet test, leukopenia with or without warning signs; the warning signs include, abdominal pain or tenderness, persistent vomiting, clinical fluid accumulation, mucosal bleeding, lethargy, restlessness, liver enlargement >2 cm, increase in HCT concurrent with rapid decrease in platelet count) within 4 days from fever onset who tested positive for onsite NS1 rapid test (SD Bio) were recruited for the study from the North Colombo Teaching Hospital, Ragama, Sri Lanka from 2015 to 2017 with informed consent. 2.5 mL blood was collected in to EDTA tubes and approximately 500 μL saliva was collected by spit method from all patients at the time of admission within 4 days from fever onset. The samples were transported and processed at 4 °C within an hour from sample collection. Isolated peripheral blood cells (PBC), plasma and saliva samples (after adding 30 mM NaOH) were stored at – 80 °C until sample analysis. A questionnaire was used to collect information pertaining to alcohol consumption, smoking habits and dietary intake. None of the patients were pregnant. Patients who later developed SD were determined according to the WHO 2009 guidelines based on the evidence of plasma leakage (pleural effusions and ascites detected using a portable bedside ultrasonogram) [32]. Routine blood tests for full blood counts and biochemical tests were carried out every day during the course of hospitalization after admission. None of the patients presented signs of SD at the time of admission within 4 days from fever onset. Twenty patients presented with signs of SD after admission during the course of infection. Samples collected from these patients at the time of admission within 4 days from fever onset were selected as the severe cases for this retrospective nested case-control study. Twenty PBC, plasma, and saliva samples collected at the time of admission within 4 days from fever onset from patients who did not present with signs of SD during the course of infection were randomly selected as controls with DF. No mortality was recorded for the patients recruited for the study.

### 2.2 Ethical statement

Ethical clearance for patient sample collection was obtained from the ethics review committee of the Faculty of Medicine, University of Kelaniya, Kelaniya, Sri Lanka (Reference number-P/119/07/2015). Samples were collected after obtaining informed written consent from all subjects. All subjects voluntarily participated in the study. All methods were carried out in accordance with the protocols approved by the ethics review committee with minimal risk to the study subjects. Patient data and personal information were stored securely. Privacy of personal information was ensured by limiting access to authorized personnel. Clinical data and patient samples were recorded with a serial study number to maintain confidentiality.

### 2.3 Quantitative Real-time PCR

Gene specific human primers against the reference gene GAPDH and iNOS were mined from previously published literature [33]. Total RNA was isolated from PBC using miRNeasy serum/plasma kit (Qiagen) with 700 μL QIAzol Lysis buffer and 140 μL chloroform according to the manufacturer’s instructions. cDNA was synthesized using the miScript II RT Kit with Hiflex buffer from 12 μL of extracted total RNA according to product manual (Qiagen). Expression of mRNA was quantified using QuantiTect SYBR Green PCR kit (Qiagen) according to product manual at an annealing temperature of 60 °C. Each reaction was carried out in triplicates in 20 μL reaction volume using StepOne real-time PCR Thermal Cycler (Applied Bio). The efficiency of amplification for iNOS was 106 % and GAPDH was 110 % based on the standard curve analysis. No-template reactions and melting curve analysis was used to confirm specificity of target amplification.

Expression levels of iNOS relative to the expression level of the reference gene in DF and SD patients were calculated as 2^-ΔCq^. Relative expression is shown as log_2_ values. Continuous variables were expressed as median and interquartile range (IQ_25-75_). The fold change of expression was calculated using the equation 2^-ΔΔCq^ and presented as log_2_ values. A difference in expression based on fold change, less than 0.5 between DF and SD cases was considered as downregulation and above 1.5 was considered as upregulation.

### 2.4. Quantification of plasma and salivary nitric oxide by Greiss reaction

Nitrite content in plasma was measured according to a previously reported method using Griess reaction against a standard series of NaNO2 [34]. Plasma sample (60 μL) was deproteinized with 7.5 μL of 200 mM ZnSO4 prior to assay for nitrites. Greiss reaction mixture was incubated at room temperature for 20 mins and absorbance at 540 nm was measured using Multiskan Go spectrophotometer (Thermo Scientific). Conversion of nitrates to nitrites by VCl3 followed by colorimetric analysis for nitrites using Greiss reaction did not give higher nitrite reading indicating that there are no detectable nitrates in the samples. Same procedure was followed to measure salivary NO.

### 2.5. Quantification of plasma and salivary oxLDL by ELISA

oxLDL content in plasma and saliva was measured using human oxLDL ELISA kit (Elabscience) according to the manufactures protocol with minor modifications. 5 μL of plasma was diluted 1:1000 in phosphate buffered saline (PBS, pH 7) and 25 μL of diluted sample was assayed. 10 μL saliva was assayed in antibody pre coated well of micro ELISA plate after dilution with 15 μL of PBS (pH 7). The oxLDL concentration was calculated based on the concentration series of reference standards of oxLDL provided with the assay kit as follows. Diluted saliva sample was removed from the well and 100 μL of biotinylated detection antibody was added into the well and incubated for 60 mins at 37 °C. Liquid was aspirated and the well was washed 3 times with wash buffer. Then 100 μL of horse radish peroxidase conjugate was added into the well and incubated for 30 mins at 37°C. Liquid was aspirated and 90 μL of substrate reagent was added washed 5 times with wash buffer and incubated for 15 mins at 37 °C. Then 50 μL of stop solution was added and the absorbance was read at 450 nm by using Multiskan Go spectrophotometer (Thermo Scientific).

### 2.6. Statistical Analysis

q-q plots and Shapiro-Wilk test were used to determine normality at a 95% confidence interval. For the Shapiro-Wilk test P > 0.05 was determined as normal distribution. Statistical significance for differentially expressed targets were determined based on the standard error of mean (SEM) of ΔCq using independent t-test. Statistically significant differences among the mean ± SD was determined using independent t-test. Statistically significant differences among the median ± median absolute deviation (MAD) was determined using Mann-Whitney U test for non-parametric independent samples. P < 0.05 was considered statistically significant. Logistic regression analysis for odds ratio, receiver operator characteristics, area under curve, specificity and sensitivity were determined using IBM SPSS Statistics, 2013 version at a 95 % confidence interval.

## 3. Results

### 3.1. Clinical characteristics at admission

Samples were collected at admission from patients recruited on, day 2 (n_DF_ = 2, n_SD_ = 3), day 3 (n_DF_= 6, n_SD_ = 12) and day 4 (n_DF_ = 12, n_SD_ = 5) from fever onset. Therefore, there were 8 DF and 15 SD samples collected within 3 days from fever onset. Most of the subjects were male (77 %), with a median age of 30 (18-60) while the female subjects had a median age of 24 (19-60) years. Participant flow diagram of the study is given in the Figure 1. At admission, there were no statistically significant differences in median laboratory clinical parameters such as thrombocytopenia, leukopenia, hematocrit count and AST and ALT levels in patients who later developed SD compared with the randomly selected DF patients included in the analysis (Table 1). These findings are consistent with the clinical parameters at admission for all patients who tested positive for dengue NS1 antigen (Supplementary Table 1). Therefore, it was not possible to make prognosis of SD based on clinical characteristics at the time of admission within 4 days from fever onset.

**FIGURE 1:**
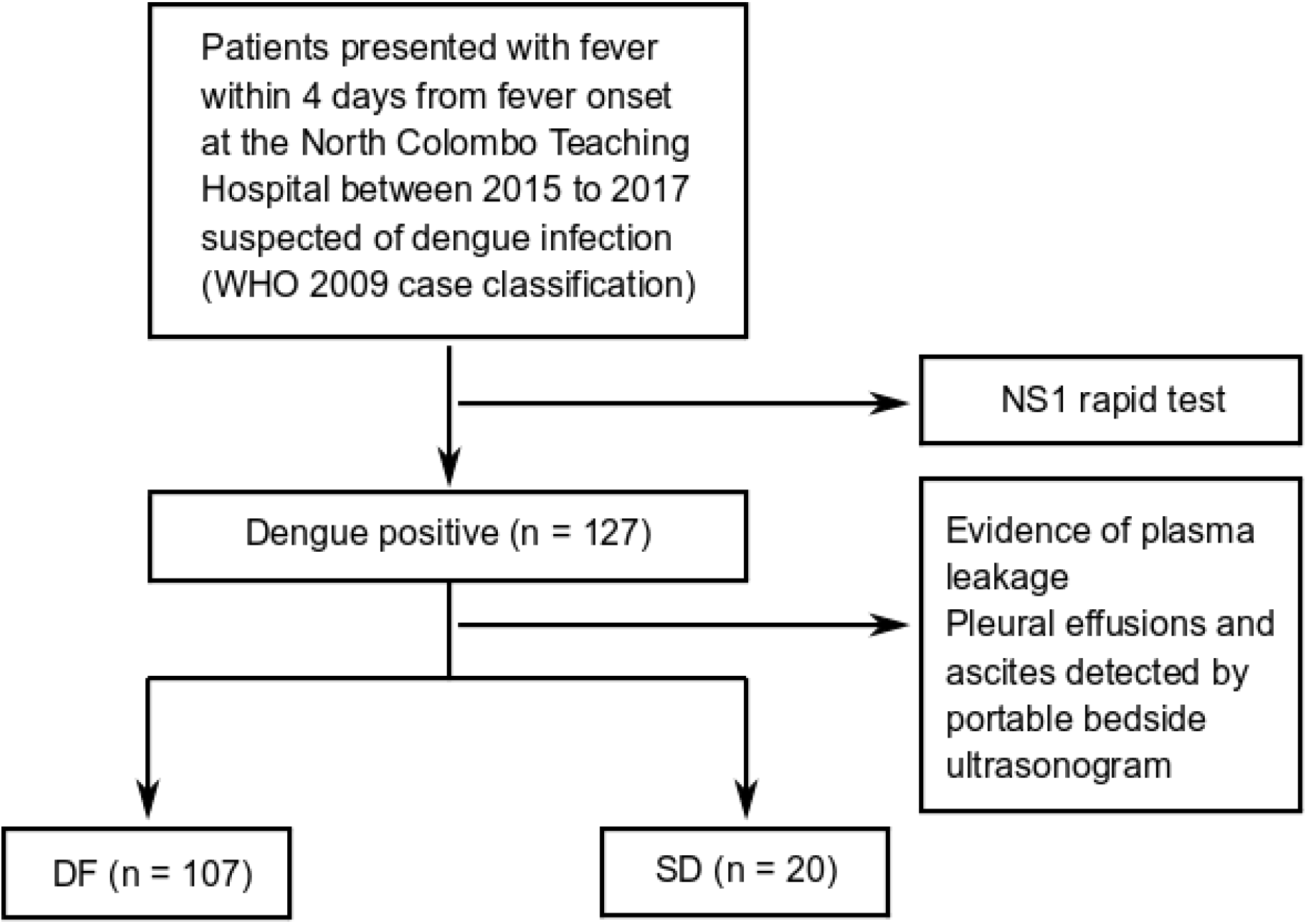
Participant flow diagram

**TABLE 1:** Clinical characteristics of dengue patients and samples collected for analysis at the admission. (Median ± MAD)

|  | DF patients<br>(n=20) | SD patients<br>(n=20) | P values |
| --- | --- | --- | --- |
| Gender (Male %/ Female %) | 68/32 | 85/15 |  |
| Age (years) | 34 (18-60) | 29 (19-60) |  |
| Platelet (X1000 cells/mm<sup>3</sup>) | 113.0±51.8 | 125.0±37.2 | 0.57 |
| Hematocrit (%) | 39.7±4.6 | 39.7± 4.0 | 0.98 |
| Hemoglobin (g/dL) | 13.7±2.5 | 13.0± 2.8 | 0.31 |
| White blood cells (X1000 cells/mm<sup>3</sup>) | 3.5±40.1 | 4.6± 60.2 | 0.39 |
| Neutrophils (%) | 64.1±18.9 | 74.5± 19.9 | 0.19 |
| Lymphocytes (%) | 27.60±12.8 | 16.0± 13.7 | 0.08 |
| Eosinophils (%) | 0.6±6.1 | 1.0± 2.2 | 0.88 |
| AST (U/L) | 34.0±39.7 | 56.0± 29.0 | 0.29 |
| ALT (U/L) | 37.1±31.1 | 43.0± 24.2 | 0.43 |

### 3.2. iNOS expression in dengue patients within four days from fever onset

Since iNOS has been implicated in host response to dengue virus infection, the level of iNOS expression was analyzed in PBC collected at the time of admission within 4 days of fever onset. The data were determined to be normally distributed. SD patients showed significantly (P < 0.05) lower iNOS expression compared to the DF patients within 4 days from fever onset. Furthermore, iNOS expression in SD patients admitted on day 3 from fever onset was also significantly (P<0.05) low compared to that of DF patients (Figure 2, Supplementary Figure 1).

**FIGURE 2:**
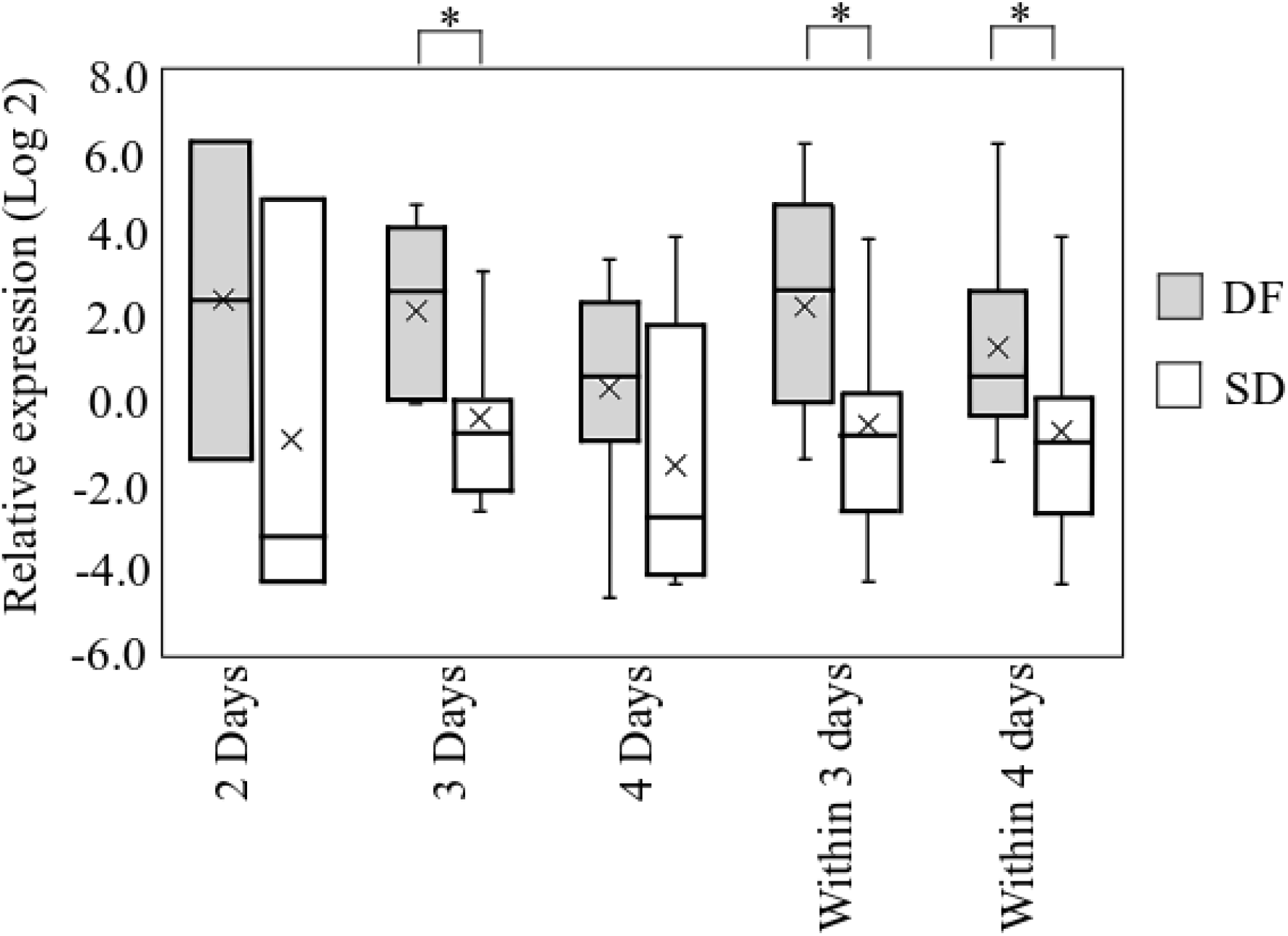
Expression of iNOS in PBC from DF patients and patients who later developed SD. Relative expression of iNOS in patient samples collected at admission from patients recruited on, day 2, day 3, day 4, within 3 days and within 4 days from fever onset designated as DF and SD. Relative expression based on 2^-ΔCq^ values against GAPDH presented as log values to the base 2. *P < 0.05 based on ΔCq ± SEM using independent t - test.

Logistic regression analysis of iNOS expression within 3 days from fever onset is predictive of SD (odds ratio, 1.74; 95 % CI 1.02 – 2.98; P < 0.05) with an area under the receiver operating curve of 0.77. The sensitivity and specificity for the development of SD were 0.73 and 0.86 respectively at ΔCq for iNOS expression of −0.09.

### 3.3. Level of nitric oxide in plasma and saliva from acute dengue patients

Expression of iNOS results in NO biosynthesis. We quantified the levels of NO in plasma using the Greiss reaction. Preliminary measurement of NO by Greiss reaction after conversion of nitrate to nitrite revealed that there was no detectable level of nitrate in the plasma samples. Therefore, the nitrite levels as measured by the Greiss reaction was taken as the total NO in the plasma samples. Median plasma NO concentration at admission within 4 days from fever onset in patients who later developed SD (17.9 ± 1.6 μM) is significantly (P < 0.05) lower than that of DF group (23.0 ± 2.1 μM). A significant decrease in median NO levels (P < 0.05) in the SD patients was observed in samples collected at admission on day 2, day 3, day 4 and within 3 days from fever onset (Figure 3). The data are not normally distributed. Therefore, Mann-Whitney U test was used to determine significant differences.

**FIGURE 3:**
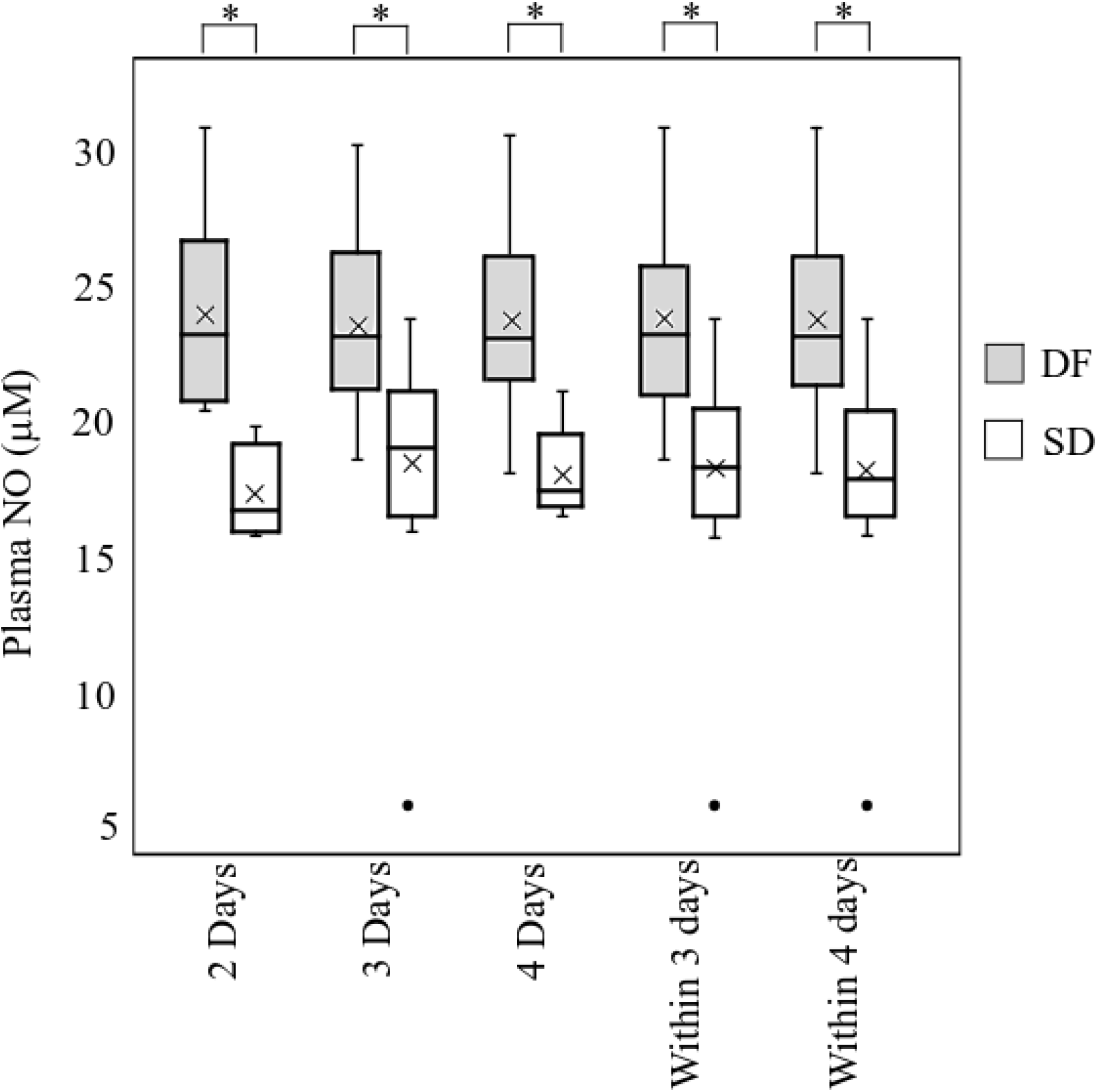
NO in plasma collected at admission within 4 days of fever onset from DF patients and patients who later developed SD. Plasma NO levels in patient samples collected on day 2, day 3, day 4, within 3 days and within 4 days from fever onset. Data expressed as median and interquartile range (IQ_25–75_). * P < 0.05

Logistic regression analysis of plasma NO level within 4 days from fever onset was found to be predictive of SD (odds ratio, 0.54; 95% CI 0.40-0.72; P < 0.05) with an area under the receiver operating curve of 0.89. The sensitivity and specificity for the development of SD within 4 days from fever onset were 0.90 and 0.70 respectively at plasma NO level of 21.3 μM (P < 0.05). The plasma NO level on day 3 and within 3 days from fever onset is also predictive of SD (odds ratio, 0.53; 95% CI 0.30-0.94; P < 0.05) with an area under the receiver operating curve of 0.87, sensitivity of 0.95 and specificity of 0.72 at plasma NO level of 22.2 μM (P < 0.05) and (odds ratio, 0.49; 95% CI 0.30-0.81; P < 0.05) with an area under the receiver operating curve of 0.90, sensitivity of 0.96 and specificity of 0.61 at plasma NO level of 22.3 μM (P < 0.05) respectively. Salivary NO levels among the patients fluctuate in a wide range in both study groups. Therefore, no significant differences were observed between the groups.

### 3.4. Oxidized LDL levels in plasma and saliva

L-arginine-NO pathway is involved in the effects of ox-LDL on platelet function [35]. Therefore, plasma oxLDL levels were analyzed at admission within four days from fever onset using ELISA. Median plasma oxLDL concentration at admission within 4 days from fever onset in patients who later developed SD (509.4 ± 179.7 ng/mL) is lower than that of DF patients at admission (688.8 ± 213.4 ng/mL). However, this difference was not statistically significant. The median oxLDL levels in the SD group (509.4 ± 224.1 ng/mL) showed a significantly (P < 0.05) low oxLDL levels in plasma collected within 3 days from fever onset compared to patients who did not develop SD (740.0 ± 300.0 ng/mL). Although the sample numbers in each group were low, the oxLDL levels in the SD group were also lower in samples collected at admission on day 2 from fever onset and significantly lower (P<0.05) on day 3 (SD = 451.9 ± 115.3 ng/mL; DF = 783.1 ± 249.4 ng/mL (Figure 4 (a)).

**FIGURE 4:**
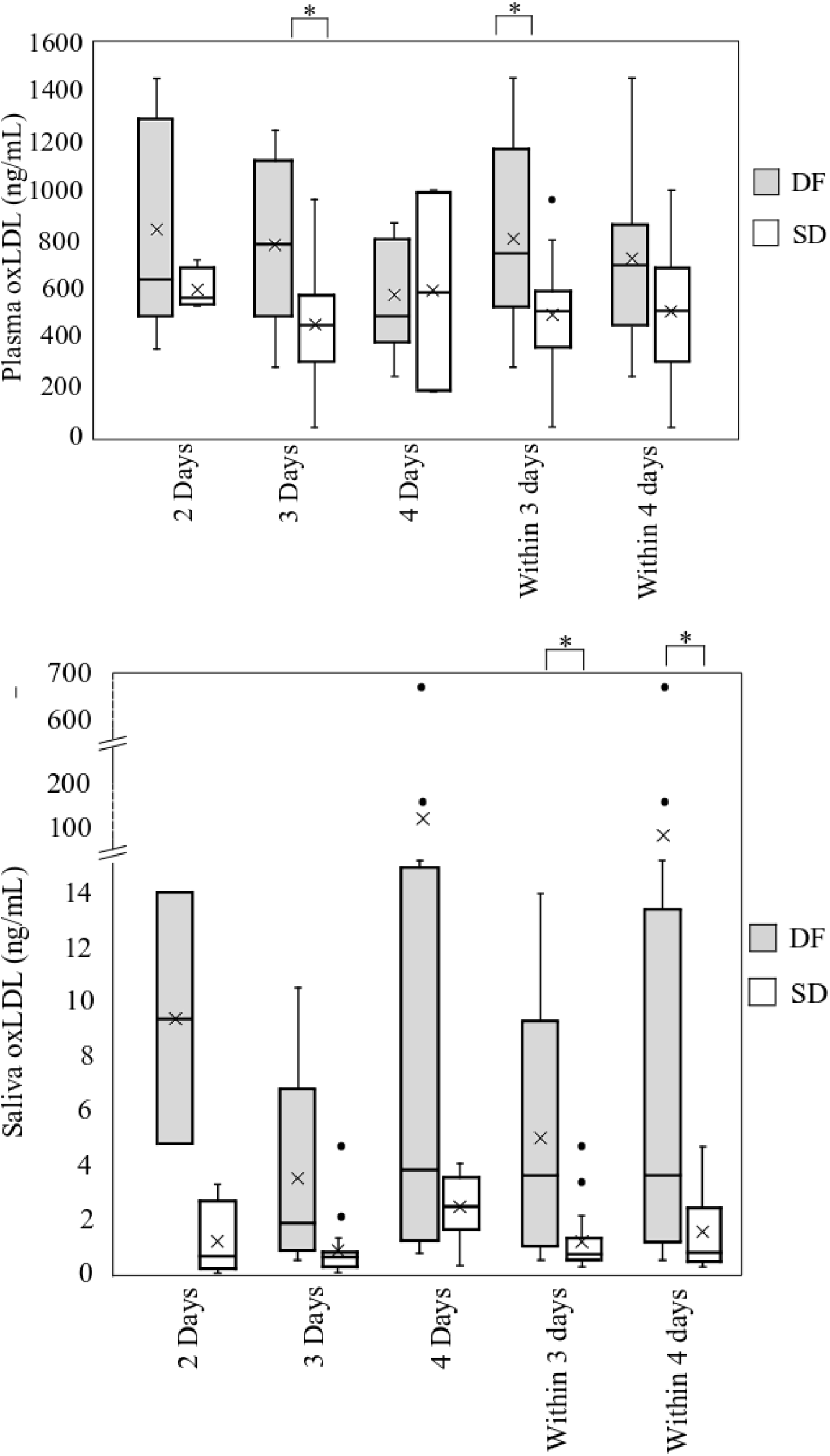
oxLDL in plasma and saliva from DF patients and patients who later developed SD collected at admission within 4 days from fever onset. oxLDL levels in (a) plasma and (b) saliva collected at admission on day 2, day 3, day 4, within 3 days and within 4 days from fever onset. Data expressed as median and interquartile range (I_Q25-75_). *P < 0.05

The adjusted odds ratio, as calculated by logistic regression analysis of plasma oxLDL level after adjustment for number of days from fever onset, was not predictive of SD within 3 days from fever onset (0.99; 95% CI 0.99-1.00, P < 0.05). However, at plasma oxLDL cut off value of 603.8 ng/mL within 3 days from fever onset, the area under the receiver operating curve is 0.76. The sensitivity and specificity for the development of SD are 0.81 and 0.77 respectively (P< 0.05).

oxLDL levels in saliva samples collected at admission within 4 days from fever onset from patients with positive diagnosis for dengue were analyzed using ELISA to evaluate whether the differences observed in the plasma samples were detectable in saliva samples as well. Median salivary oxLDL in SD patients at admission, within 4 days from fever onset (0.8 ± 0.5 ng/mL) is significantly lower (P < 0.05) than that of DF patients (3.6 ± 2.6 ng/mL). The median salivary oxLDL at admission, within three days from fever onset in SD patients (0.7 ± 0.3 ng/mL) is also significantly lower (P < 0.05) than that of DF patients (3.6 ± 2.6 ng/mL) suggesting that salivary oxLDL may also serve as an early non-invasive marker for SD (Figure 4 (b)).

Logistic regression analysis of salivary oxLDL levels at admission within 4 days from fever onset was found to be predictive of SD (odds ratio, 0.69; 95% CI 0.49-0.97, P <0.05) with an area under the receiver operating curve of 0.75. The sensitivity and specificity for the development of SD were 0.57 and 0.91 respectively at salivary oxLDL level of 0.9 ng/mL, 0.67 and 0.58 respectively at salivary oxLDL level of 2.0 ng/mL, 0.76 and 0.55 respectively at salivary oxLDL level of 2.3 ng/mL and 1.00 and 0.50 respectively at salivary oxLDL level of 4.6 ng/mL (P < 0.05).

## 4. Discussion

iNOS expression, plasma NO, plasma oxLDL and salivary ox LDL levels show significant differences between the DF patients and patients who later developed SD during the acute phase of infection. iNOS is implicated in host response to dengue virus infection and plasma leakage symptomatic of SD [25]. iNOS has been reported to overexpress in response to dengue infection with a corresponding increase in the NO levels compared to healthy subjects (25). In the present study, patients who developed severe symptoms show significantly (P<0.05) lower levels of iNOS expression and a corresponding decrease in the NO levels compared to those who did not develop severe symptoms during the early phase of infection. This may be due to the role of iNOS and NO as part of the host defense mechanism which appears to be compromised even during the early phase of infection in the individuals who later developed SD.

Due to the role of iNOS in NO biosynthesis, corresponding decrease in NO levels were expected as a result of observed downregulation of iNOS expression in SD patients during the acute phase of infection, which was significant within 2 days from fever onset and remained significantly low in plasma during the acute phase of infection. These findings are consistent with the reported differences of serum NO and nitrite levels suggesting their role as early marker of disease severity for dengue [29]. However, the plasma NO levels are at least four fold higher than serum NO levels with a clear distinction of the levels between DF patients and patients who later developed SD with an odds ratio of approximately 0.5 for samples collected on day 3, day 4, within 3 days and 4 days from fever onset, indicating that plasma NO may serve as a more robust marker.

Oxidation of lipids by NO can result in increased levels of plasma oxLDL which promotes vasoconstriction and platelet activation with alterations in platelet function which has been connected to dengue-associated plasma leakage [35, 36]. Low plasma oxLDL levels corresponding to the low plasma NO was observed at admission within 4 days from fever onset in dengue patients who later developed SD. Therefore, the plasma oxLDL levels do not appear to participate in development of symptoms of SD such as plasma leakage during the early phase of infection.

Similarly, salivary oxLDL levels showed a significant decrease in the patients who later developed SD compared to the DF patients within 4 days of infection proving to be an excellent non-invasive biological source for predictive markers for dengue. Salivary oxLDL levels have been shown to correlate with the serum oxLDL levels [31]. oxLDL has also been associated with cardiovascular risks [31]. Therefore, oxLDL levels in a larger cohort of dengue patients during acute phase of infection may be necessary to validate the role of oxLDL in saliva as a non-invasive early prognostic biomarker of SD and evaluate the effect of the confounding factors. Although saliva may serve as a non-invasive source for NO levels, saliva was proven to be an unreliable biological source due to high standard deviation of NO concentration that may be resulting from the influence of oral health and diet [37].

Our study is limited by the relatively small sample sizes for 2, 3 and 4 days from fever onset and relatively few samples from female patients to assess the potential of these markers to predict the outcome within these parameters. Therefore, further analysis in a larger cohort within each day from fever onset during the acute phase is needed to assess the full potential of these biomarkers to distinguish DF patients from those who progress to SD during the early stages of infection. We were unable to determine whether dietary habits, social habits such as smoking and non-communicable conditions such as high cholesterol and diabetes influence the iNOS expression, plasma NO and plasma and salivary oxLDL among the DF and SD patients within four days from fever onset, due to unreliable response rate from the subjects.

## 5. Conclusions

Differential expression of iNOS, plasma NO and salivary oxLDL levels may serve as reliable early biomarkers to predict the development of SD within 4 days from fever onset. Our findings also suggest saliva as a potential new non-invasive biological source for early prognosis of disease outcome for dengue.

## Data Availability

The data used to support the findings of this study are included within the article and the supplementary files.

## Supplementary Materials

Supplementary Table 1: Clinical characteristics of dengue fever and severe dengue fever patients at the admission. Supplementary Figure 1: Fold change of iNOS expression in PBC between DF and SD patients at admission recruited on, day 2, day 3, day 4, within 3 days and within 4 days from fever onset.

